# QuimP – Analyzing transmembrane signalling in highly deformable cells

**DOI:** 10.1101/171199

**Authors:** Piotr Baniukiewicz, Sharon Collier, Till Bretschneider

## Abstract

**Summary:** Transmembrane signalling plays important physiological roles, with G protein–coupled cell surface receptors being particularly important therapeutic targets. Fluorescent proteins are widely used to study signalling, but the analysis of image time series can be challenging, in particular when changes in cell shape are involved. To this end we have developed QuimP software. QuimP semi-automatically tracks cell outlines, quantifies spatio-temporal patterns of fluorescence at the cell membrane, and tracks local shape deformations. QuimP is particularly useful for studying cell motility, for example in immune or cancer cells.

**Availability and Implementation:** QuimP (http://warwick.ac.uk/quimp) consists of a set of Java plugins for Fiji/ImageJ (http://fiji.sc/) and can be easily installed through the Fiji Updater (http://warwick.ac.uk/quimp/wiki-pages/installation). It is compatible with Mac, Windows and Unix-based operating systems, requiring version >1.45 of Fiji/ImageJ and Java 8. QuimP is released as open source (https://github.com/CellDynamics/QuimP/) under an academic licence.

**Supplementary Information:** Supplementary materials (SI-A to SI-D) are available at Bioinformatics online. Test data is available from http://warwick.ac.uk/quimp/test_data.

## 1. Introduction

In transmembrane signalling the cell membrane plays a fundamental role in localising intracellular signalling components to specific sites of action, for example to reorganise the actomyosin cortex during cell polarisation and locomotion. The localisation of different components can be directly or indirectly visualised using fluorescence microscopy, for high-throughput screening commonly in 2D. A quantitative understanding demands segmentation and tracking of whole cells and fluorescence signals associated with the moving cell boundary, for example those associated with actin polymerisation at the cell front of locomoting cells. Different approaches are reviewed in (Barry *et al.*, 2015) and (Ryan *et al.*, 2013). As regards segmentation, a wide range of methods can be used (threshold based, region growing, active contours or level sets) to obtain closed cell contours, which then are used to sample fluorescence adjacent to the cell edge in a straightforward manner. The most critical step however is cell edge tracking, which links points on contours at time t to corresponding points at t+1. Optical flow methods have been employed, but usually fail to meet the requirement that total fluorescence must not change. QuimP uses a method (ECMM, electrostatic contour migration method (Tyson *et al.*, 2010) which has been shown to outperform traditional level set methods. ECMM minimises the sum of path lengths connecting all pairs of points, equivalent to minimising the energy required for cell deformation. The original segmentation based on an active contour method and outline tracking algorithms have been described in (Dormann *et al.*, 2002; Tyson *et al.*, 2010; Tyson *et al.*, 2014).

(Fig. 1) outlines a typical QuimP workflow for analysing cell motility.

**Fig. 1.**
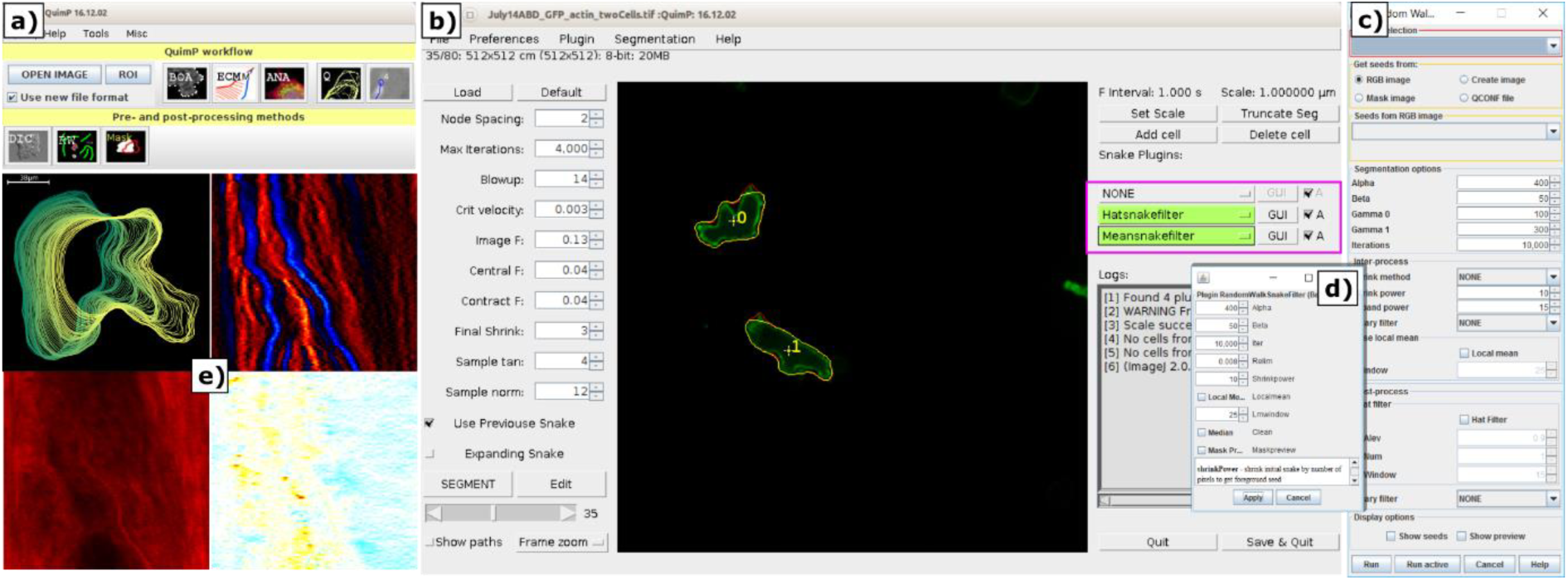
**(a)** QuimP toolbar, with tools arranged in the order of a typical workflow. Upper row: Open image time series, and main data analysis plugins (BOA: cell segmentation, ECMM: contour tracking, ANA: sampling of cortical fluorescence, QA: detailed quantitative analysis and visualisation in the form of spatial-temporal maps, PA: protrusion analysis (experimental, working Matlab routines are provided)). Bottom row: Pre- and post-processing plugins (DIC: DIC image reconstruction, RW: customised random walk segmentation, Mask: Cell outline to mask converter). **(b)** BOA segmentation window. Purple frame highlights novel support of external contour filters. **(c)** Interface for the new random walk segmentation module. (d) New BOA plugin that integrates RW and AC methods. **(e)** Exemplary results from the QA module (clockwise): cell outlines, convexity map, cell boundary displacement map, fluorescence map.

## 2. New features

Here we describe a completely revised version of QuimP, which is being released as source code for the first time (current version 2017-07-01). A user manual is available from the QuimP homepage. Main new features including walkthrough examples are described in detail in Supplementary Information SI-A. These are 1) use of the JSON file format to store the complete analysis workflow and facilitate exchange of data with software like Matlab (SI-B). Being able to save and restore the complete workflow is particularly useful when segmenting long sequences that require manual corrections. 2) Three new modules have been added, for reconstructing differential interference contrast (DIC) images, customised random walk cell segmentation, and generation of image masks from segmented cell outlines. 3) A new architecture supports writing custom vector filters that can directly operate on cell contour data, without requiring deep knowledge of QuimP itself; currently available are a running mean filter, protrusion removal filter (SI-A), and a bridge filter that allows to call the random walk module directly from BOA. Masks generated by any other segmentation method available in ImageJ can be used as input for further QuimP analysis (SI-A). 4) Improved segmentation by combining QuimP’s original active contour segmentation with a modified random walk method (Fig. 1c): Active contour methods are a good choice for segmenting cells, but notoriously struggle when dealing with highly concave cell outlines. The original random walk method (Grady, 2006) is superior in this respect, but has difficulties with strong gradients in fluorescence along the cell axis, as typically observed for many proteins involved in cell polarisation in directed cell movement. We have implemented a locally adaptive version in QuimP which overcomes this problem.

The random walk method is a supervised learning method that requires users to label (seed) image pixels that belong to foreground (cell) and background. However, we can employ masks obtained by a preliminary active contour segmentation to seed these class labels for each frame of a time series automatically (Details and figures are presented in SI-A and SI-C). Foreground and background pixels are assigned after contracting and expanding masks using ECMM (Tyson *et al.*, 2010). ECMM preserves the shape of the contour and prevents that thin cellular processes are eroded (SI-D), significantly improving the segmentation of cellular protrusions and cavities. We evaluated the quality of the new method using typical time-series of migrating cells tagged with fluorescent markers for different cytoskeletal proteins. 750 image frames were manually segmented (gold standard), and compared to results achieved with QuimP run in an unsupervised manner, i.e. without changing parameters between frames. The modified random walk method significantly reduces the Hausdorff distance (maximum distance between the segmented contour and the gold standard) and the number of false positive pixels per contour length (SI-C and SI-D).

## 3. Conclusions

The strength of the built-in method for contour tracking (ECMM) and its user friendliness have gained QuimP some recognition already, resulting in more than 70 publications to date where it has been used (http://warwick.ac.uk/quimp/quimp-refs/). The recent developments described here, namely making QuimP available as open source, with frequent updates and extended documentation, improvements in segmentation quality, and important changes to the architecture to support customised cell contour filters, are important steps on the way to turn QuimP into a sustainable resource.

## Acknowledgements

### Funding

This work is supported by BBSRC Bioinformatics and Biological Resources grant BB/M01150X/1.

### Conflict of Interest

none declared.

